# Metformin Impairs Intestinal Fructose Metabolism

**DOI:** 10.1101/2023.04.17.537251

**Authors:** Wenxin Tong, Sarah A. Hannou, Ashot Sargsyan, Guo-Fang Zhang, Paul A. Grimsrud, Inna Astapova, Mark A. Herman

## Abstract

**Objective:** To investigate the effects of metformin on intestinal carbohydrate metabolism *in vivo*.

Method: Male mice preconditioned with a high-fat, high-sucrose diet were treated orally with metformin or a control solution for two weeks. Fructose metabolism, glucose production from fructose, and production of other fructose-derived metabolites were assessed using stably labeled fructose as a tracer.

**Results:** Metformin treatment decreased intestinal glucose levels and reduced incorporation of fructose-derived metabolites into glucose. This was associated with decreased intestinal fructose metabolism as indicated by decreased enterocyte F1P levels and diminished labeling of fructose-derived metabolites. Metformin also reduced fructose delivery to the liver. Proteomic analysis revealed that metformin coordinately down-regulated proteins involved carbohydrate metabolism including those involved in fructolysis and glucose production within intestinal tissue.

**Conclusion:** Metformin reduces intestinal fructose metabolism, and this is associated with broad-based changes in intestinal enzyme and protein levels involved in sugar metabolism indicating that metformin’s effects on sugar metabolism are pleiotropic.

**Highlights:**

- Metformin decreases intestinal fructose absorption, metabolism, and fructose delivery to the liver.
- Metformin reduces intestinal glucose production from fructose-derived metabolites.
- Metformin reduces protein levels of multiple metabolic enzymes involved in fructose and glucose metabolism in intestinal tissue.

## Introduction

Metformin is a first-line anti-diabetic drug and the most widely prescribed oral anti-hyperglycemic worldwide [1]. Beneficial effects of metformin have also been suggested for people with obesity [2], non-alcohol fatty liver disease [3], Alzheimer’s disease [4], and some cancers [5; 6]. Even though metformin has been investigated intensively for decades, the target organ(s) and the underlying mechanism of metformin’s beneficial effects remain controversial.

Metformin’s anti-diabetic effects are often attributed to its ability to reduce endogenous glucose production [7; 8]. Because the liver is the major site of gluconeogenesis in mammals, most studies have focused on metformin’s effects on liver tissue [9]. However, the role of liver in metformin’s actions is receiving more scrutiny as therapeutic doses of metformin may not produce sufficient concentrations of metformin in hepatocytes to mediate its observed systemic effects [8; 10]. In contrast with liver, metformin is highly concentrated in intestinal tissue [11; 12]. In intestinal tissue, metformin may achieve concentrations that are 100 to 300-fold higher than circulating plasma levels, and ~ 2-10-fold higher than that in the liver [11; 12]. This suggests that metformin has the potential to have outsized impact on intestinal function to regulate systemic glucose homeostasis [13]. Metformin may regulate glycemia through effects on the intestine via multiple mechanisms. For instance, a recent study showed that metformin’s glucose-lowering effect is mediated through activating the PEN2-directed lysosomal AMPK pathway in intestinal tissue and not in liver [14]. Additionally, multiple studies in both humans and preclinical models using a F18-fluorodeoxyglucose (F18-FDG) tracer indicate that metformin enhances glucose uptake in the colon and possibly the small intestine [13; 15; 16]. Recent studies from our group and others demonstrated that fructose is metabolized at high rates in intestinal tissue and contributes to intestinal glucose production [17; 18]. In this study, using fructose to assess enterocyte metabolic function and as a gluconeogenic tracer, we show that short-term treatment of mice with metformin impairs intestinal carbohydrate metabolism and reduces intestinal glucose production adding to the mechanisms by which metformin impacts systemic glucose homeostasis.

## Methods

### Materials

Metformin (hydrochloride) was purchased from Cayman Chemical (No. 13118). D-Fructose was purchased from VWR international (VWRV0226). [U-^13^C]-fructose, and [U-^13^C]-fructose-6-phosphate (M6-F6P) were purchased from Cambridge Isotope Labs (CLM-1553-PK, and CLM-8616 respectively). Norvaline and D-[1,6-^13^C_2_]-fructose (M2-fructose) were purchased from Sigma-Aldrich (53721 and 587613 respectively).

### Animals

All animal experiments were approved by Duke Institutional Animal Care and Use Committee. 17-18 weeks old wild-type C57/BL6J (Stock# 000664) male mice were purchased from The Jackson Laboratory and maintained in a pathogen free vivarium at Duke University. All animals were maintained at ~ 22°C on a 12-hour light-dark cycle (6 AM to 6 PM) and adapted for at least one week before any experimental procedure or intervention. Within experiments, mice were divided randomly into either water control or metformin treatment groups. To prepare the metformin oral-gavage solution, 500 mg metformin was dissolved in 20 mL sterile water and aliquoted into 14 vials stored at 4 °C. After pretreatment of all mice with 1 week of high-fat, high-sucrose diet (45% fat, 20% sucrose, Research Diet, D12451i), we randomly distributed mice to two groups and orally gavaged water or metformin solutions at a dose of 10 µL/g body weight. After two weeks of metformin or water administration, an i.p. glucose tolerance test (GTT) was conducted. Two days after the i.p. GTT was performed, mice were sacrificed 30 minutes after fructose gavage (1:1 mixture of unlabeled fructose and [U-^13^C]-fructose). Unless otherwise noted, all procedures including oral gavage and body weight measurement were performed at 9:30 AM. Euthanasia after fructose gavage was performed at 2 PM, 5 hours after food removal at 9 AM.

### Glucose tolerance test

Animals were fasted for 5 hours starting at 9 AM. At 2PM, animal body weight was measured, and sterile glucose solution was injected intraperitoneally at a dose of 1.5 g/Kg body weight. Glycemia was measured using a drop of blood drawn from a nicked tail vein via glucometer (Bayer Contour) at 0, 15, 30, 60, 90, and 120 minutes after injection.

### Animal euthanasia and sample/tissue harvest

Animals were euthanized under isoflurane anesthesia 30 minutes after fructose gavage (1:1 mixture of unlabeled fructose and [U-^13^C]-fructose at 4g/Kg body weight). Mice were anesthetized under continuous oxygen following the guidelines recommended by the Duke Institutional Animal Care and Use Committee. Once deep anesthesia was confirmed by a toe pinch, blood samples from the tail vein were harvested. Laparotomy was subsequently performed, and the portal vein was exposed. Clean scissors were used to make a small incision in the portal vein, and blood (200 µL) was extracted using a 1 mL syringe from the peritoneal cavity and transferred to an EDTA-coated blood collection tube (Sarstedt, 16.440.100). The liver and intestine were removed from the abdominal cavity. Liver tissue was snap frozen in liquid nitrogen. The small intestine extending from 0.5 cm distal to the end of the stomach and extending to 0.5 cm proximal to the cecum was excised, flushed with ice-cold PBS and snap frozen in liquid nitrogen.

### Gas chromatography-mass spectrometry (GC/MS)

Plasma was prepared from collected blood by centrifugation at 2,000 g, 4°C for 15 minutes. Tissue samples were powdered under liquid N_2_. 0.2 mM 2-Deoxy-D-glucose (2DG), 0.2 mM M2-fructose, 0.1 nM M6-F6P or 0.1 mmol norvaline was added to 10 μL plasma samples or 10 mg powdered tissue samples as an internal standard for the measurement of glucose, fructose or TCA cycle intermediates respectively. After that, 400 μL MeOH, 400 μL ddH2O, and 400 μL chloroform were added sequentially and briefly vortexed after each addition. After centrifugation at 12,000 g for 15 minutes, supernatants were collected and dried under nitrogen.

Relative quantification and enrichment measurements for glucose, glycerate, F1P, and GA3P: Dried sample residues were derivatized with methoxylamine hydrochloride and trimethylsilyl (TMS) sequentially. Specifically, 25 μL of methoxylamine hydrochloride (2% (w/v) in pyridine) was added to the dried residues and incubated for 90 minutes at 40°C before the addition of 35 μL of TMS and incubation for 30 minutes at 60°C. The samples were then centrifuged for 2 minutes at 12,000 g and the supernatants of derivatized samples were transferred to GC vials for further analysis.

Absolute quantification of fructose: the dried residues were derivatized with methoxylamine hydrochloride and propionic anhydride sequentially adapted from a previously described method [19]. 25 μL of methoxylamine hydrochloride (2% (w/v) in pyridine) was added to the dried residues and incubated for 60 minutes at 70°C. After centrifugation for 2 minutes at 12,000 g, 50 μL propionic anhydride was added and incubated for 30 minutes at 60°C prior to another round of drying under nitrogen. The dried residues were resuspended with 55 μl pure ethyl acetate and transferred to GC vials for analysis.

Relative quantification and enrichment measurements of TCA cycle intermediates: Dried residues were derivatized with methoxylamine hydrochloride and tertbutyldimetheylchloros (TBDMS) sequentially as previously described [3]. 25 μL of methoxylamine hydrochloride (2% (w/v) in pyridine) was added to the dried residues and incubated for 90 minutes at 40°C before the addition of 35 μL of TBDMS and incubation for 30 minutes at 60°C. The samples were then centrifuged for 2 minutes at 12,000 g and the supernatants of derivatized samples were transferred to GC vials for further analysis. GC/MS analysis was conducted using an Agilent 7890B GC system and Agilent 5977A Mass Spectrometer. 1 μL of the derivatized sample was injected into the GC column. The GC temperature gradient started at 90°C, increased to 260°C at the speed of 9°C per minute and further increased to 290 °C at the speed of 30°C per minute. Then the temperature was held at 290°C for 5 minutes with a run time of 24.9 minutes. Ionization was conducted by EI at 70 eV with Helium flow at 1 mL/min. Temperatures of the source, the MS quad, the interface, and the inlet were maintained at 230°C, 150°C, 280°C, and 250°C, respectively. Mass spectra were recorded in mass scan mode from m/z 50 to 700. The levels of metabolites were calculated based on the area of the peak at its specified retention time. Total ion counts (A.U.) of metabolites were assessed as the sum of its isotopologues and normalized to their specific standards as detailed above and below.

### Stable Isotope Analysis and Average Carbon-13 Label Incorporation Rates

Isotopologues containing 0, 1, to n heavy atom(s) in a molecule were referred as M+0, M+1, to M+n respectively. The retention times of individual metabolites and their isotopomers are described in Supplementary Table 1. Isotope enrichment and labeling analysis in this study were corrected for natural isotope distribution as previously described [20; 21]. Specifically, the stable isotope enrichment of each metabolite in plasma and tissue samples was corrected based on the natural isotope distribution measured in the same type of sample treated without labeled molecules. For example, the natural isotope distribution matrix of individually measured metabolites in the blood samples were experimentally determined by assaying and averaging the blood samples from animals who had not received any treatment with isotopically labeled sources.

The average carbon-13 incorporation was calculated according to the equation as previously described in reference (18). Specifically, n_M+i_ refers to the number of ^13^C carbons in the isotopomer (M+i). For example, n = 0 for M+0, n = 1 for M+1, and n = 2 for M+2, and so on. “A” refers to the corresponding percentage abundance of the isotopomer. N refers to the total carbon number, which for glucose, N = 6.

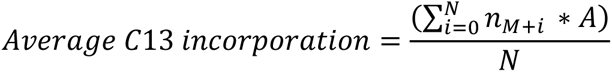

### Quantification of metabolites

The relative quantification of total glucose, F1P, and GA3P levels were calculated as the ratio derived from the sum of total ion counts of all isotopologues to their specific internal standards (2DG for glucose and M6-F6P for F1P and GA3P respectively). The absolute quantification of M+0 and M+6 fructose isotopologues were calculated as the ratio of M+0 fructose ion counts or M+6 fructose ion counts to M+2 fructose ion counts which is derived from the M2-fructose internal standard respectively.

For the quantification of glucose derived from any fructose carbons, we first calculated the stable isotope enrichment of each isotopologue of glucose after correction for the natural isotope distribution. Because fructose was administered as an equal mixture of unlabeled and [U-^13^C]-labeled fructose, levels of glucose derived from any fructose carbons were calculated as twice the total glucose ion counts multiplied by the corrected percentage of the sum of glucose isotopomers containing any ^13^C. Levels of glucose with three fructose-derived carbons were calculated as twice the total glucose ion counts multiplied by the corrected percentage of M+3 glucose isotopomers.

### Proteomics Sample Preparation

Samples from mice treated with and without metformin (n=5) underwent large-scale analysis of relative protein abundance changes as described previously [22]. For each sample, approximately 15 mg of pulverized mouse intestinal tissue was solubilized in ice-cold urea Lysis buffer (8 M urea in 50 mM Tris, pH 8.0, 40 mM NaCl, 2 mM MgCl_2_, 1x Roche cOmplete ULTRA EDTA-free protease inhibitor mini tablet, 1x PhosStop phosphatase inhibitor tablet, 5 mM NaF and 10 mM Na pyrophosphate). Samples were lysed with a combination of disruption with a TissueLyzer (Qiagen) for 30 sec at 30 Hz, subjected to three freeze-thaw cycles, and sonication with a probe sonicator. After centrifugation at 10,000 g for 10 min at 4°C, protein concentration in the supernatant was determined by BCA, and a 400 μg aliquot of each was adjusted to 2 mg/mL with Urea Lysis Buffer. To reduce and alkylate cysteine residues, samples were incubated with 5 mM DTT at 32 °C for 30 min, cooled to RT, and incubated with 15 mM iodoacetamide for 30 min in the dark at RT. Following quenching of unreacted iodoacetamide by the addition of equimolar DTT (up to 15 mM), samples were enzymatically digested with LysC (100:1 w/w, protein to enzyme) at 32 °C for 4 hr. After dilution to 1.5 M urea with 50 mM Tris (pH 8.0), 5 mM CaCl_2_, samples were digested with trypsin (50:1 w/w, protein:enzyme) overnight at 32 °C. Samples were acidified to 0.5% TFA and centrifuged at 10,000 ×g for 10 min at RT to pellet insoluble material. Supernatant containing soluble peptides was subjected to solid phase extraction (SPE) with a Waters 50 mg tC18 SEP-PAK column to remove salt, eluting once with 500 μL 25% acetonitrile/0.1% TFA and twice with 500 μL 50% acetonitrile/0.1% TFA. The eluate was frozen and dried by speed vac. Samples were resuspended in 100 μL of 200 mM triethylammonium bicarbonate (TEAB), vortexed, mixed with a different 10-plex Tandem Mass Tag (TMT) reagent (0.8 mg resuspended in 50 μL 100% acetonitrile), shaken for 4 hr at RT, and quenched with 0.8 μL 50% hydroxylamine (shaking for 15 min at RT). All samples were mixed, frozen, and dried by speed vac. The mixture was resuspended in 1 mL 0.5% TFA and subjected to SPE again with a Waters 100 mg tC18 SEP-PAK SPE column as described above and the eluate was frozen and dried by speed vac. The 4 mg TMT-labeled peptide mixture was fractionated by high pH reversed-phase (HPRP) chromatography. After each of 12 fractions was lyophilized and resuspended in 1 mL 80% acetonitrile/0.15% TFA, 50 μL was removed for analysis of unmodified peptides, frozen, and dried by speed vac.

### Proteomics data acquisition

Samples were analyzed by nanoflow liquid chromatography (nLC) on an EASY-nLC 1200 UH-PLC, followed by electrospray ionization (ESI) with an EASY-Spray source, and tandem mass spectrometry (MS/MS) with a Q Exactive Plus Hybrid Quadrupole-Orbitrap (Thermo Fisher Scientific). Sample fractions were resuspended in 22 μL of 0.1% FA and were analyzed with triplicate runs, injecting 1 μL for the first and adjusting the volume for subsequent runs to target an MS1 base peak intensity of 1 ×10^10^. For each injection, the sample was first trapped on an Acclaim PepMap 100 C18 trapping column (3 μg particle size, 75 μm ×20 mm) with solvent A (0.1% FA) at a variable flow rate dictated by a maximum pressure of 500 bar, after which the analytical separation was performed over a 140 min gradient (flow rate of 300 nL/min) of 2% to 40% solvent B (90% ACN, 0.1% FA) using an Acclaim Pep-Map RSLC C18 analytical column (2 μg particle size, 75μg ×500 mm column (Thermo Fisher Scientific) with a column temperature of 55 °C. MS1 (precursor ions) was performed at 70,000 resolution with an AGC target of 3 ×10^6^ ions and a maximum injection time (IT) of 60 ms. MS2 spectra (product ions) were collected by data-dependent acquisition of the top 10 most abundant precursor ions with a charge greater than 1 per MS1 scan, with dynamic exclusion enabled for a window of 30 s. Precursor ions were filtered with a 0.7 m/z isolation window and fragmented with a normalized collision energy of 30. MS2 scans were performed at 35,000 resolution, with an AGC target of 1 ×10^5^ ions and a maximum IT of 60 ms. All raw files were uploaded to Proteome Xchange (PXD041833) and jPOST (JPST002144) [23; 24]

### Proteomics data analysis

Raw files were processed with Proteome Discoverer 2.5 (PD2.5) from Thermo Fisher Scientific. A complete mouse proteome FASTA file containing 55,315 reviewed (Swiss-Prot) and unreviewed (TrEMBL) protein sequences, downloaded from UniProt on 8/15/2022, was imported into PD2.5. “Treatment” (metformin) versus “Water” (control) were incorporated as “Study Factors”. MS1 precursor mass recalibration was performed after a pre-search using the Spectrum Files RC node. Subsequent searches with Sequest HT were conducted at 10 ppm (recalibrated) and 0.02 Da mass tolerances for precursor and fragment ions, respectively. Modifications considered included oxidation on methionine (variable + 15.995 Da), carbamidomethylation of cysteine (fixed + 57.021 Da), and TMT6plex addition to lysine peptide N-termini (fixed + 229.163 Da). Searches allowed for up to 2 missed K/R cleavages by trypsin (full specificity). Percolator [9] was used to filter peptide spectral matches (PSMs) to an estimated FDR of less than 1%. The Peptide Isoform Grouper node was used to collapse PSMs down to unique peptides, while maintaining FDR < 1% at the peptide level. The Protein FDR Validator node was used to group peptides into proteins using the rules of strict parsimony and maintain FDR < 1% at the protein level. The Reporter Ion Quantifier node was used to filter PSMs for quantitative quality, setting an upper threshold 50% precursor co-isolation and an average reporter ion S/N> 2.5. For PSMs meeting these criteria, quantitative values were summed together at the peptide and protein levels (after adjusting for TMT isotope impurities). Unique peptides and Razor peptides were used for protein quantitation calculations, excluding peptides with oxidized methionine. Quantitative values were normalized for any deviations from equal total peptide signal on each TMT “channel”. Data were further analyzed in R. PCA analysis was performed within each conditions, water and metformin specifically, for outlier detection. One sample in each group appeared to be an extreme outlier compared to the other samples in the group and were not used in subsequent analysis. Data analysis was conducted in R 4.0. Differential expression analysis was conducted with the Limma and DEseq2 packages [25; 26]. Data were visualized with the ggplot2 package and hierarchical clustering heatmaps were generated using the pheatmap package where Pearson correlation was used as the distance measure.

### Western Blot

20 to 30 mg intestinal tissue samples were homogenized in 0.5 mL cell signaling lysis buffer that contains 20 mM Tris-HCl, 150 mM NaCl, 1 mM Na2EDTA, 1 mM EGTA, 1% Triton and further supplemented with protease inhibitors (Sigma-Aldrich, P8340) and phosphatase inhibitor cocktail (Pierce, A32957). After centrifugation at 12,000 g for 15 minutes, the supernatant was transferred to a fresh tube to measure protein concentration using the BCA method (ThermoFisher, 23225). Approximately 35 ug of protein was loaded per lane for SDS-PAGE electrophoresis. After electrophoresis, immunoproteins were transferred to PVDF membranes. The membranes were then incubated with primary antibodies overnight at 4 °C with gentle rocking after blocking with 5% solution of BSA (Cytivia, SH30574.02) at room temperature for 1 hour. Primary antibodies used in this work are as follows: anti-AMPK (Cell Signaling, 5831S), anti-AMPK (phospho T172) antibody (Cell Signaling, 2635S), and anti-P85 (Upstate, 06-496). Blots were incubated with appropriate secondary antibody after washing with TBST for 3 times. Chemiluminescence was conducted using the Clarity Western ECL substrate kit (Bio-Rad, 1705061) and membranes were imaged on ChemiDoc XP (Bio-Rad). Quantification of blots was conducted using ChemiDoc XP associated Image Lab software v6.0. For loading normalization, whole lane proteins were quantified using Bio-Rad Stain free technology.

### Statistics

All data are presented as mean ± SEM. Comparison of metabolite levels between the metformin group and controls were conducted using a two-tailed student’s t-test in GraphPad Prism. Statistical significance was assumed at p less than 0.05.

## Results

### Short-term metformin treatment improves glucose tolerance without altering fasting glycemia

To investigate metformin’s role in intestinal metabolism, 18-week-old wildtype C57BL/6J mice were preconditioned with 1 week of high-fat, high-sucrose diet (45% Fat, 40% Sucrose) and then orally gavaged with metformin at a daily dose of 250 mg/kg of body weight for two weeks. This dose of metformin in mice is reported to achieve circulating concentrations of ~ 5 - 10 µM which is comparable to that in humans with Type-2 Diabetes (T2D) taking 1 - 2 g metformin per day [27].

Metformin treatment did not affect body weight (Figure 1A, metformin: 33.5 ± 2.7 vs water: 34.2 ± 2.1 g). This is similar to observations that short-term metformin treatment does not significantly impact body weight in human patients [28]. Fasting glycemia remained similar between the metformin and control groups (Figure 1B). Whereas metformin had no significant effect on body weight or fasting glycemia, 2 weeks of metformin enhanced glucose tolerance (Figure 1C and D) with a reduction in glycemia at the 30-minute time point (metformin: 292.44 ± 92.57 mg/dL vs water: 393.89 ± 87.23 mg/dL, p<0.05) and decreased area under the curve (metformin: 27790 ± 3326 A.U. vs water: 34600 ± 2676 A.U., p<0.05). Thus, short-term metformin treatment enhanced glucose tolerance without significant changes in body weight.

**Figure 1.**
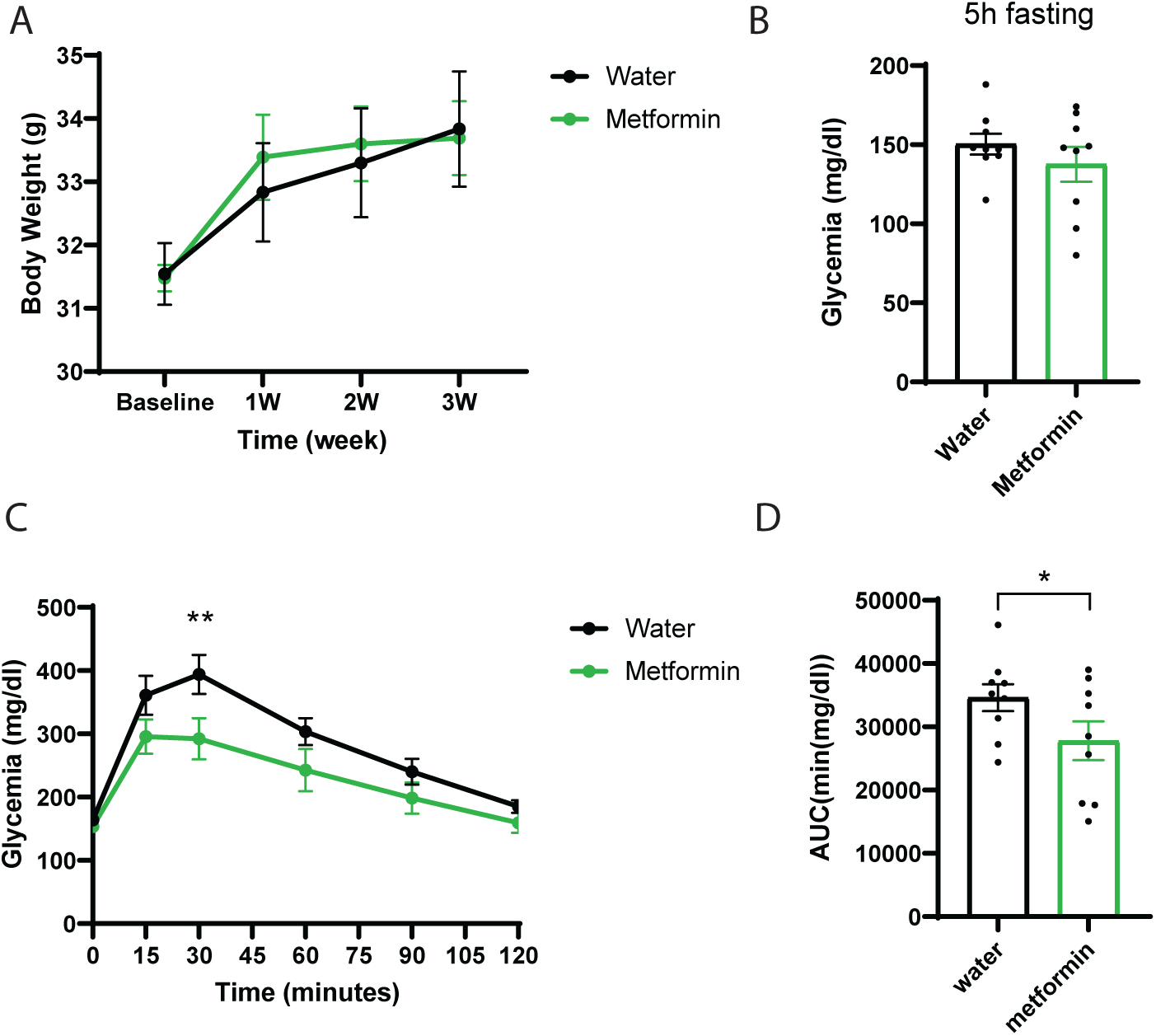
Short-term metformin treatment improves glucose tolerance without reducing body weight and fasting glycemia levels. (A) Body weight before and after metformin versus water treatment. (B) Fasting glycemia levels after 2 weeks of treatment. (C - D) Intraperitoneal glucose tolerance test after 2 weeks of treatment. Data represent means ± SEM. n=9 per group * p<0.05; ** p<0.01; *** p<0.001. Analysis performed via paired student-t test.

### Short-term metformin treatment decreases intestinal fructose metabolism and fructose delivery to the liver

The intestine is capable of robust fructose metabolism and a large portion of intestinally metabolized fructose is used to produce glucose within intestinal tissue (Figure 2A) [17; 18]. After transport into the cell, fructose is first metabolized to fructose-1-phosphate (F1P) followed by further metabolism to dihydroxyacetone phosphate (DHAP) and glyceraldehyde (GA). Glyceraldehyde can be converted into glyceraldehyde-3-phosphate (GA3P) via triokinase or into glycerate via aldehyde dehydrogenase. Glycerate production tends to increase following a bolus of fructose and saturation of triokinase [29]. Fructose-derived triose phosphates then enter the glycolytic/gluconeogenic triose-phosphate pool and can be used either for glucose production or can be further metabolized to pyruvate/lactate and potentially enter the TCA cycle for oxidation. We used ^13^C-labeled fructose as a tracer to investigate the effects of metformin on intestinal fructose and carbohydrate metabolism including glucose production. After a 5 hour fast, we gavaged animals from both metformin and control groups with a 1:1 mixture of unlabeled fructose and fructose that was universally labeled with carbon-13 ([U-^13^C]-fructose).

**Figure 2.**
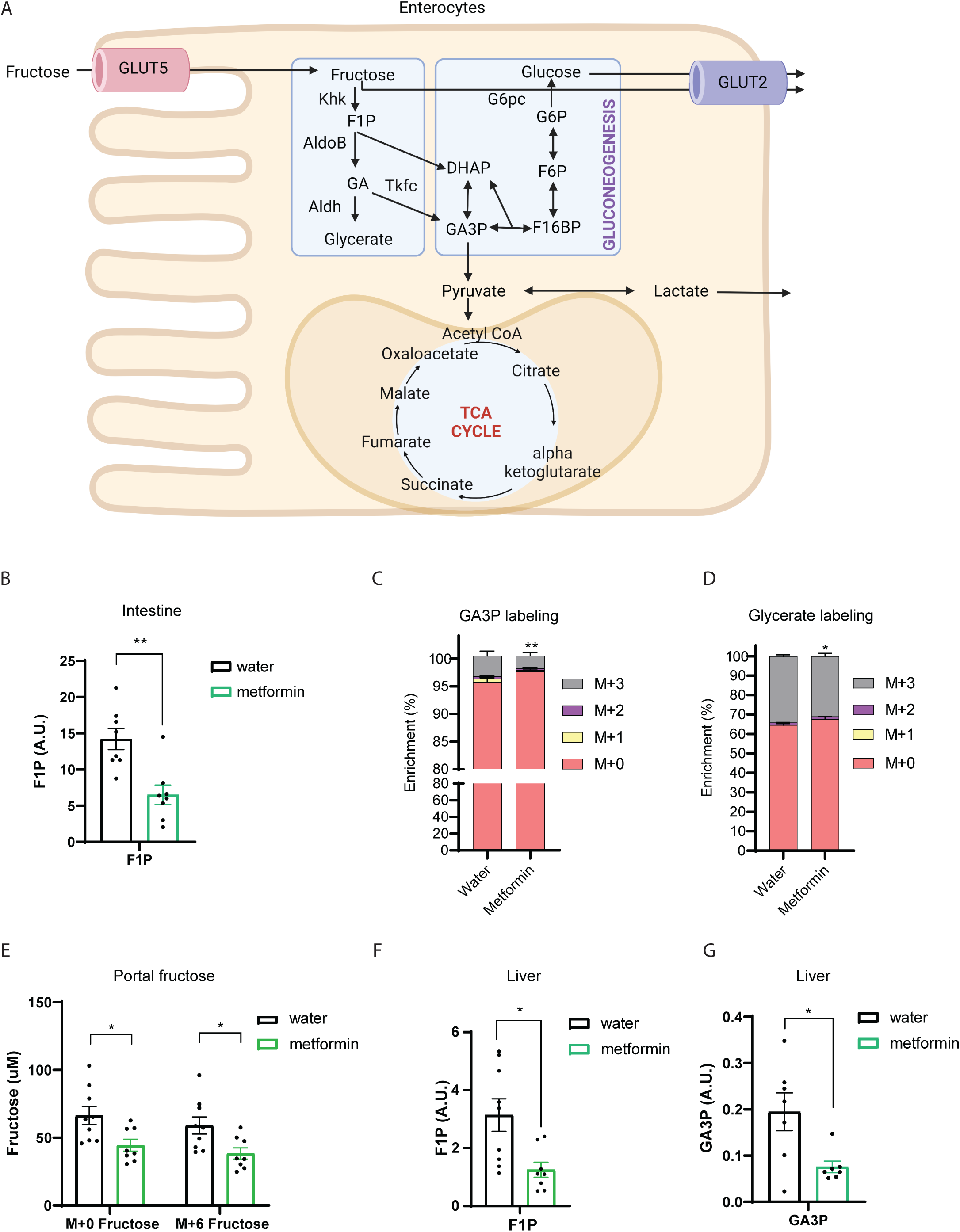
Short-term metformin treatment is sufficient to decrease fructose catabolism in the intestinal tissue and fructose delivery to the liver. (A) Diagram showing the fructose catabolism pathways within enterocytes. (B) Relative quantity of fructose 1-phosphate, and enrichment of (C) glyceraldehyde 3-phosphate, and (D) glycerate in the intestinal tissue 30 minutes after fructose gavage in mice treated with water versus metformin for 2 weeks. (E) Absolute fructose levels measured in the portal blood. (F) Relative quantification of fructose 1-phosphate and (G) glyceraldehyde 3-phosphate in the liver tissue after fructose gavage. 0.1 nmol M6-Fructose 6-phosphate and 4 nmol M2-fructose were used as internal standard for the measurement of fructose 1-phosphate and fructose respectively. Norvaline was used as the internal standard for the measurement of glyceraldehyde 3-phosphate, and glycerate. Data represent means ± SEM. * p<0.05; ** p<0.01; *** p<0.001. Analysis performed via paired student-t test.

The levels of F1P were markedly reduced (from 14.20 ± 3.85 to 6.50 ± 3.54 A.U., p<0.01) in the intestinal tissue of animals treated with metformin versus water (Figure 2B). Metformin treatment reduced the enrichment of isotope-labeled three-carbon metabolites, glyceraldehyde 3-phosphate (GA3P) and glycerate in intestinal tissue (Figure 2C and D). These data indicate that fructose catabolism in intestinal tissue is impaired by metformin treatment. Reductions in intestinal fructose metabolism could lead to increased hepatic fructose delivery and contribute to fructose-induced metabolic disease [30]. However, we observed that metformin treatment decreased portal vein M+0 and M+6 fructose levels by ~ 34% (Figure 2E). Reduced intestinal fructose metabolism in combination with decreased portal fructose levels indicates that metformin impairs intestinal fructose absorption. Additionally, the reduction in portal fructose following metformin treatment indicates that metformin may reduce fructose delivery to the liver. Consistent with this, metformin reduced levels of hepatic F1P (3.14 ± 1.59 to 1.25 ± 0.68 A.U., p<0.05, Figure 2F). Metformin treatment produced similar reductions in hepatic GA3P levels (0.19 ± 0.10 to 0.08 ± 0.03 A.U., p<0.05, Figure 2G). Altogether, these results indicate that two weeks of metformin treatment is sufficient to decrease intestinal fructose metabolism and reduce fructose delivery to the liver.

### Short-term metformin treatment blunts intestinal glucose production

At euthanasia—30 minutes after fructose gavage—metformin reduced glucose levels within intestinal tissue by 47% compared to control (Figure 3A). The reduced glucose levels within intestinal tissue could be caused either by an increase in intestinal glucose metabolism as suggested by experiments using FDG-glucose as a tracer [13], reduced intestinal glucose production, or a combination of the two. Within intestinal tissue, metformin also reduced the percentage of glucose containing fructose derived labeled carbons (Figure 3B). At this time point, glycemia in systemic circulation is only reduced by 13% in metformin treated mice compared to controls (Figure 3C). The labeling of intestinal glucose with fructose-derived, labeled carbons is ~ 2-fold higher than in peripheral circulation (Figure 3B and 3D). In combination, these results show that glucose derived from fructose is being produced locally within the intestinal tissue. Glucose produced from fructose within enterocytes is generated following metabolism of fructose to trioses and will therefore predominately contain three fructose-derived carbons (Figure 2A and 3B). Within intestinal tissue, metformin reduced the labeling of glucose compared to controls (Figure 3B) and reduced the levels of glucose containing any or three fructose-derived carbons (Figure 3E and F) to a larger extent than glucose without fructose derived carbons (Figure 3G). The reduction in intestinal glucose containing fructose-derived carbons accounts for most of the reduction in intestinal glucose levels in the metformin treated mice (comparing Fig 3A and 3E). Thus, the reduction in intestinal glucose production from fructose is a major contributor to reduced intestinal glucose levels following metformin treatment. This is consistent with metformin’s effect to reduce fructose absorption and metabolism as indicated in Figure 2.

**Figure 3.**
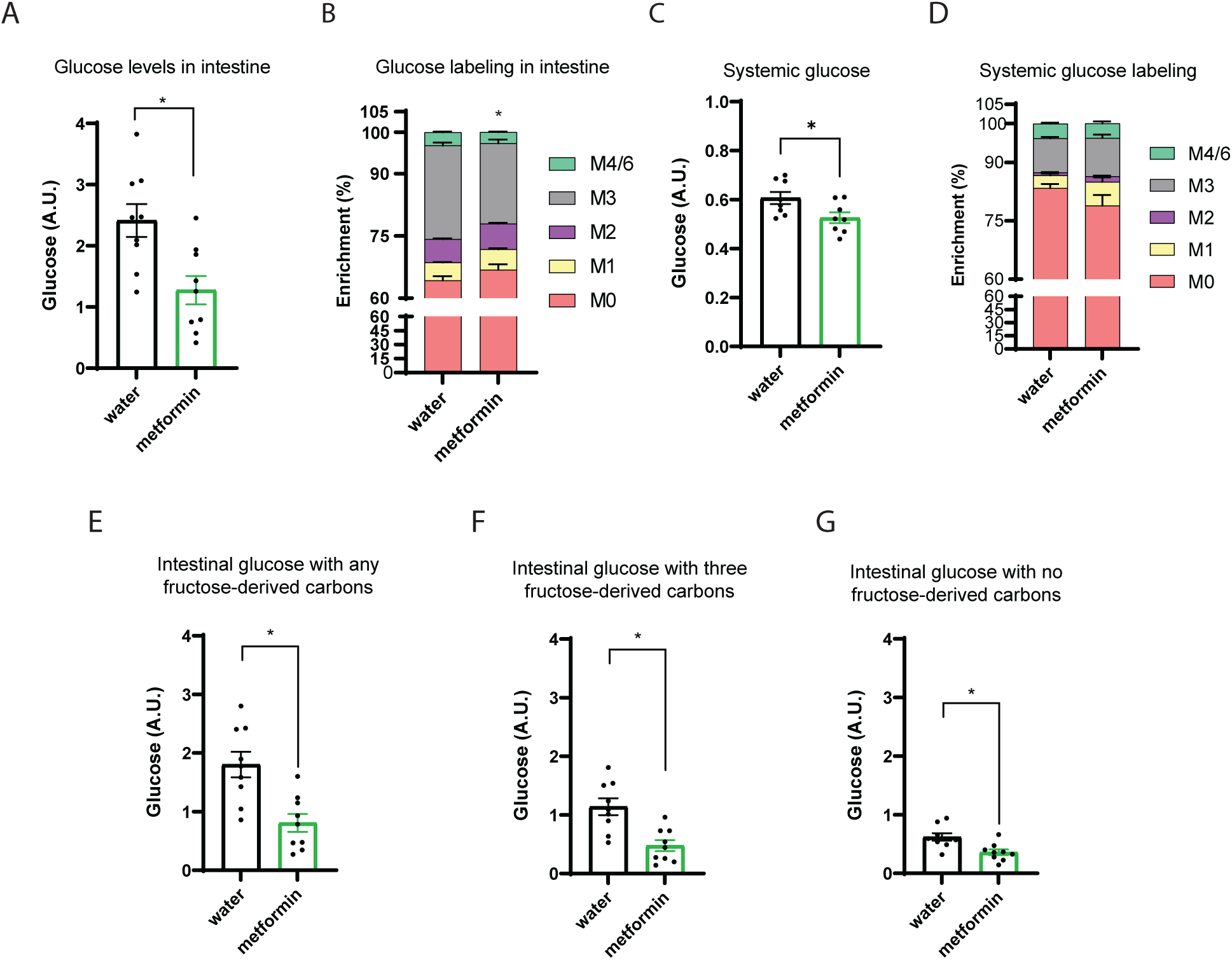
Short-term metformin treatment reduces glucose production from fructose in intestinal tissue. (A) Relative glucose levels and (B) the enrichment of glucose with labeled fructose tracer in intestinal tissue. (C) Relative circulating glucose and (D) enrichment of glucose measured in tail vein plasma. The levels of glucose with (E) any, (F) three, and (G) no carbons derived from fructose in intestinal tissue. 2 nmol 2-Deoxu-D-glucose (2DG) was used as internal standard for the measurement of glucose. Data represent means ± SEM. * p<0.05; ** p<0.01; *** p<0.001. Analysis performed via paired student-t test.

### Short-term metformin treatment decreases fructose incorporation into the TCA cycle

We previously demonstrated that fructose gavage rapidly labels the intestinal TCA cycle pool which is a major contributor to circulating TCA cycle metabolites [31]. We next investigated whether metformin also affects this process. The enrichment of labeled TCA cycle intermediates was reduced in intestinal tissue of metformin treated mice (Figure 4A-C). Metformin treatment produced a similar reduction in portal plasma TCA cycle metabolite enrichment (Figure 4D-F). These data support our previous finding that the intestine is a major source of circulating TCA cycle intermediates. Moreover, these results indicate that labeling of circulating TCA cycle metabolites is responsive to metformin treatment and measurement of this labeling pattern in blood may be a useful surrogate index of intestinal metabolic function.

**Figure 4.**
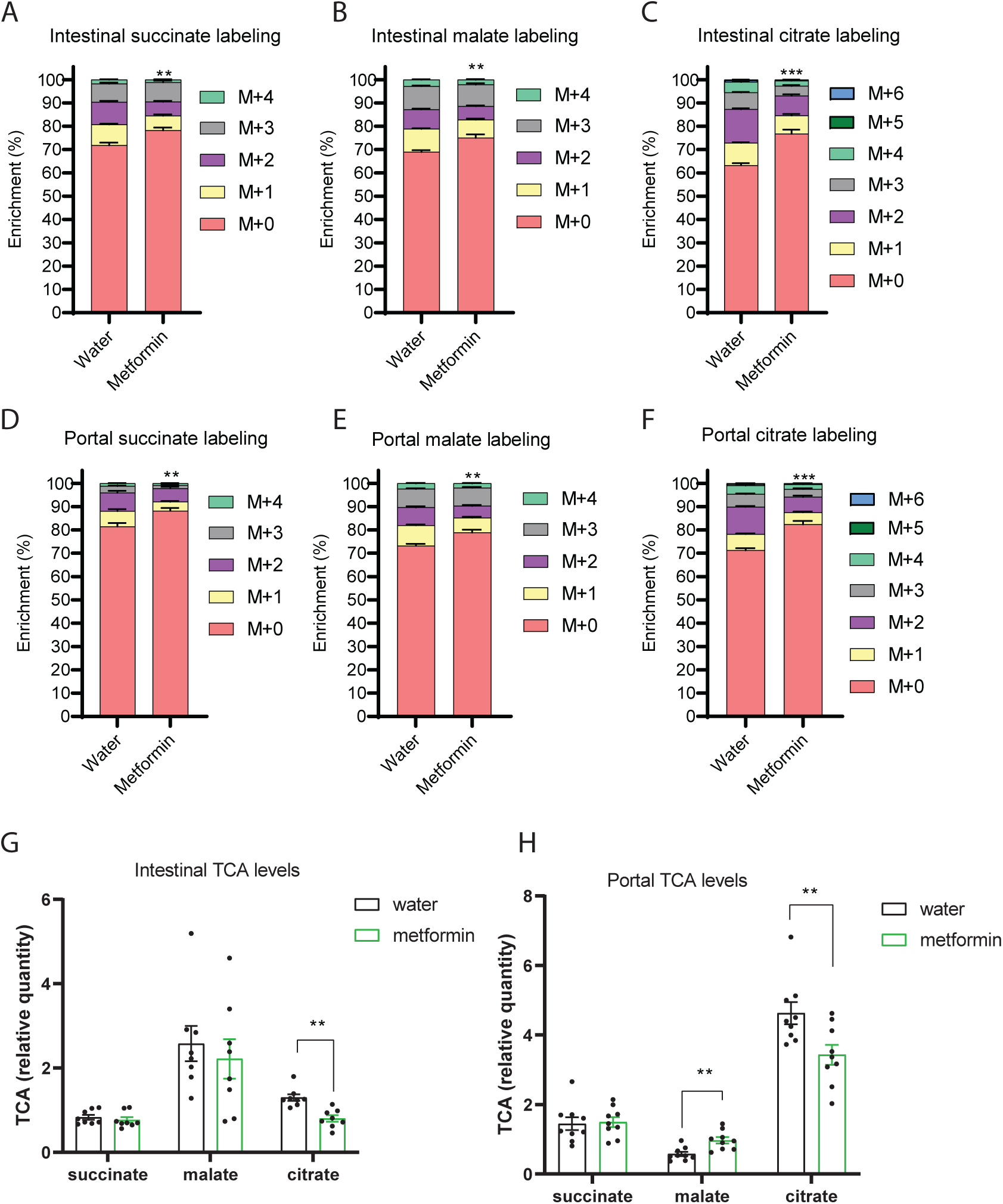
Short-term metformin treatment reduces production of intestinal TCA cycle intermediates from fructose. The enrichment of (A) succinate, (B) malate, and (C) citrate in intestinal tissue 30 minutes after fructose gavage in mice treated with water versus metformin for 2 weeks. The enrichment of (D) succinate, (E) malate, and (F) citrate in the portal plasma after fructose gavage. Relative quantification of TCA cycle metabolites in the (G) intestinal tissue, and (H) portal circulation. Norvaline was used as the internal standard for these measurements. Data represent means ± SEM. * p<0.05; ** p<0.01; *** p<0.001. Analysis performed via paired student-t test.

Whereas metformin uniformly reduced TCA cycle enrichment in both intestinal tissue and portal blood samples, its effect on TCA cycle metabolite levels was mixed. Metformin reduced citrate levels in both intestinal tissue and portal circulation (Figure 4G and H). In contrast, metformin produced no changes in either tissue or circulating succinate and produced an isolated increase in portal malate levels without a change in malate in intestinal tissue although the variance was high for malate measurements in tissue. The changes in TCA cycle metabolite levels may suggest that metformin inhibits malate to citrate conversion in the TCA cycle, potentially via inhibition of malate dehydrogenase. This would be consistent with metformin’s purported effects to inhibit Complex I activity as malate dehydrogenase donates electrons to Complex I and its activity is closely coupled to that of the electron transport chain [32–35].

### Metformin reduces the abundance of intestinal proteins involved in carbohydrate metabolism

As isotope-tracing metabolomics indicated that metformin reduced intestinal fructose catabolism and glucose production, we next aimed to investigate molecular mechanisms contributing to these changes via proteomics in intestinal tissue. In total, 5,328 proteins were quantified and annotated via our discovery proteomics (Supplementary Table 2). Principal component analysis (PCA) using all quantified proteins identified two clusters that separate the treatment groups (Figure 5A). However, the 95% confidence intervals for the metformin and control clusters overlap suggesting that global protein expression levels within the intestine were not markedly different between treatment conditions (Figure 5A).

**Figure 5.**
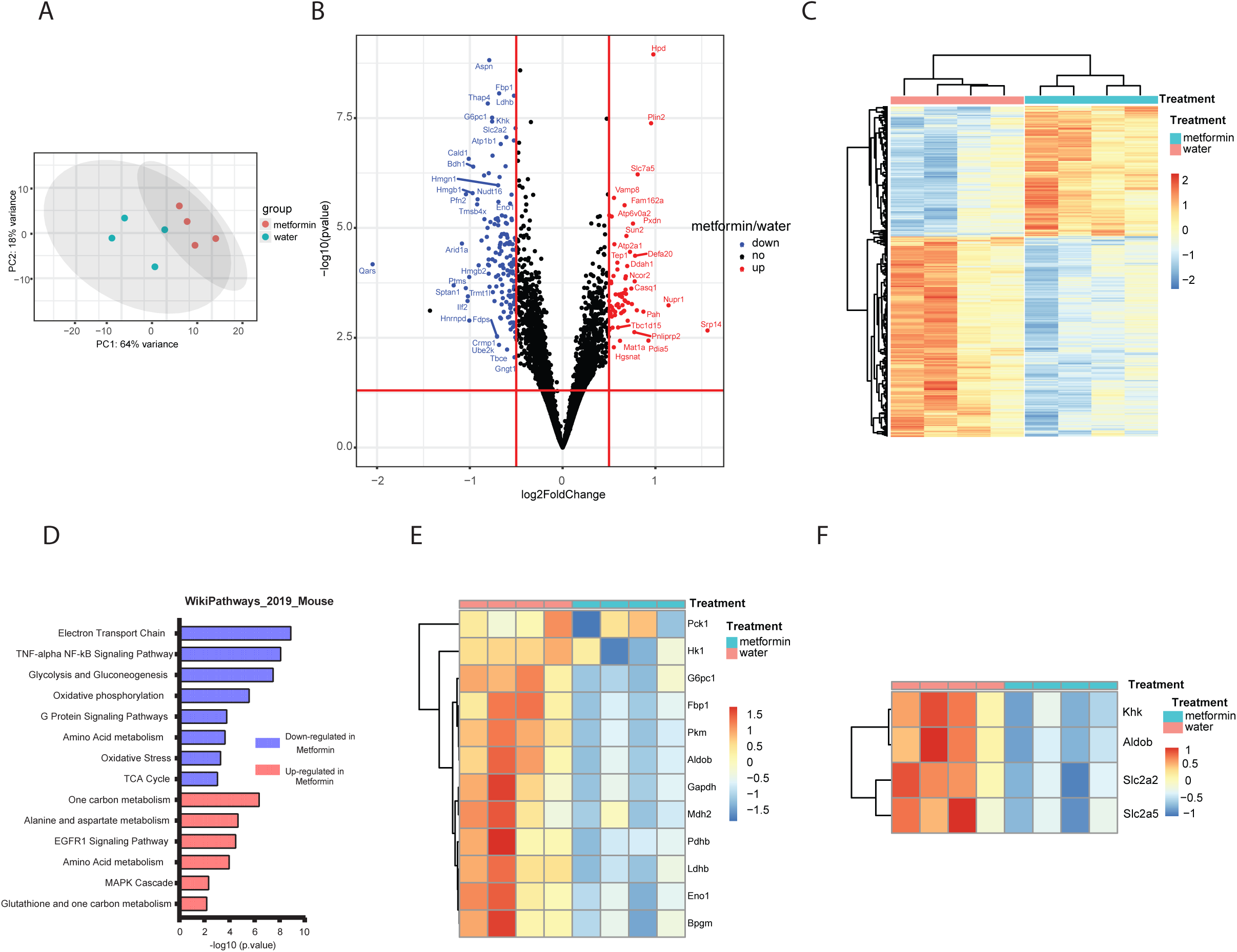
Proteomics demonstrates down-regulation of proteins involved in carbohydrate metabolism in intestinal tissue. (A) Principal component analysis of quantitative proteomics data generated from the intestinal tissue samples of mice treated with water versus metformin. (B) Volcano plot and (C) K-means clustering of differentially abundant proteins in the intestinal tissue samples from water versus metformin groups. (D) Pathway analysis using proteins that are significantly down-regulated in the intestinal tissue samples of metformin versus water group. (E) Significantly altered protein levels in the gluconeogenesis pathway as well as key hexose transporters and fructolytic enzymes. Analysis performed in R using data generated from Proteome Discoverer.

A total of 1,942 proteins were identified as differentially abundant between metformin and control groups at p < 0.05 (two-tailed student’s t-test). After adjusting for repeated measures by the Benjamini and Hochberg method, a total of 219 proteins were identified as differentially abundant at P_adjusted_ < 0.05 and |log_2_ fold-change| > 0.5 (Figure 5B and Supplementary Table 3). Many of the top-ranked differentially abundant proteins are associated with metabolic process including gluconeogenic enzymes such as Fbp1 and G6pc (Figure 5B). Despite variability in protein abundance levels within groups, unsupervised K-means clustering segregated metformin treated from untreated mice (Figure 5C). Pathway analysis using differentially expressed proteins with P_adjusted_ < 0.05 suggested that metformin treatment affected multiple metabolic pathways in the intestinal tissue (Supplementary Table 4 and 5). One of the top down-regulated pathways in intestinal tissue from the metformin group is “Glycolysis and Gluconeogenesis (WP157)” (Figure 5D). Other metabolic pathways demonstrating downregulation with metformin treatment include the “Electron Transport Chain (WP295)” and “Oxidative Phosphorylation”, pathways potentially complementary to the putative effects of metformin to inhibit mitochondrial function [36; 37] (Figure 5D). In addition, proteins associated with both glycolysis and gluconeogenic pathways, including pyruvate carboxykinase (Pck1), Fructose 1,6-bisphosphatase (Fbp1), glucose 6-phosphatase (G6pc1), lactate dehydrogenase (Ldhb), pyruvate dehydrogenase (Pdh1), malate dehydrogenase (Mdh2), enolase 1 (Eno1), hexokinase (Hk1), pyruvate kinase (Pkm), bisphosphoglycerate mutase (Bpgm), were markedly down-regulated in the metformin group compared to controls (Figure 5E). Similarly, hexose transporters capable of fructose transport including both GLUT5 (Slc2a5) and GLUT2 (Slc2a2), as well as key fructolytic enzymes ketohexokinase (Khk) and aldolase, fructose-bisphosphate B (Aldob) were all significantly down-regulated in metformin treated intestinal tissues (Figure 5F). The reduction in glycolytic, gluconeogenic, and fructolytic transporters and enzymes are consistent with the results from the metabolomic tracer studies that indicate that metformin impairs intestinal fructose absorption and catabolism as well as glucose production.

### Activation of AMPK signaling is likely to impact metformin’s effects on intestinal metabolic functions

The proteomics study suggests that mitochondria, and especially the electron transport chain, are impacted by metformin treatment through coordinate downregulation of many of its constituent proteins (Figure 5D, Figure 6A, and Supplementary Table 4). The ability of metformin to impact mitochondrial function is well documented [36–39]. Activation of AMPK is purported to play a central role in metformin’s mechanism of action and activation of AMPK may be mediated via metformin’s ability to inhibit mitochondrial Complex I and via other mechanisms [36; 40; 41]. We examined the abundance levels of detectable proteins quantified in our proteomics data that are annotated within the “AMPK signaling pathway (KEGG: mmu04152)” and conducted a PCA analysis using these 50 proteins. As is indicated in Figure 6B, abundance of components of the AMPK signaling pathway in the intestine samples from metformin versus control treated mice tend to segregate based on the first and second principal components (PC1 and PC2, Supplementary Table 5). Our intestinal proteomics study only detected the α (Prkaa1, Prkaa2) and ƴ subunits (Prkag1) of AMPK component (Figure 6C). Among these, the abundance of Prkaa1 significantly decreased after metformin treatment while Prkaa2 tended to increase (Figure 6C). To further examine the role of AMPK signaling, we measured the total protein levels of AMPKα and the phosphorylation of AMPKα at the Thr 172 site, a marker of AMPK activation. As expected, metformin treatment increased the phosphorylation level of AMPKα Thr172 indicating that short-term metformin treatment activates intestinal AMPK signaling (Figure 6D and E).

**Figure 6.**
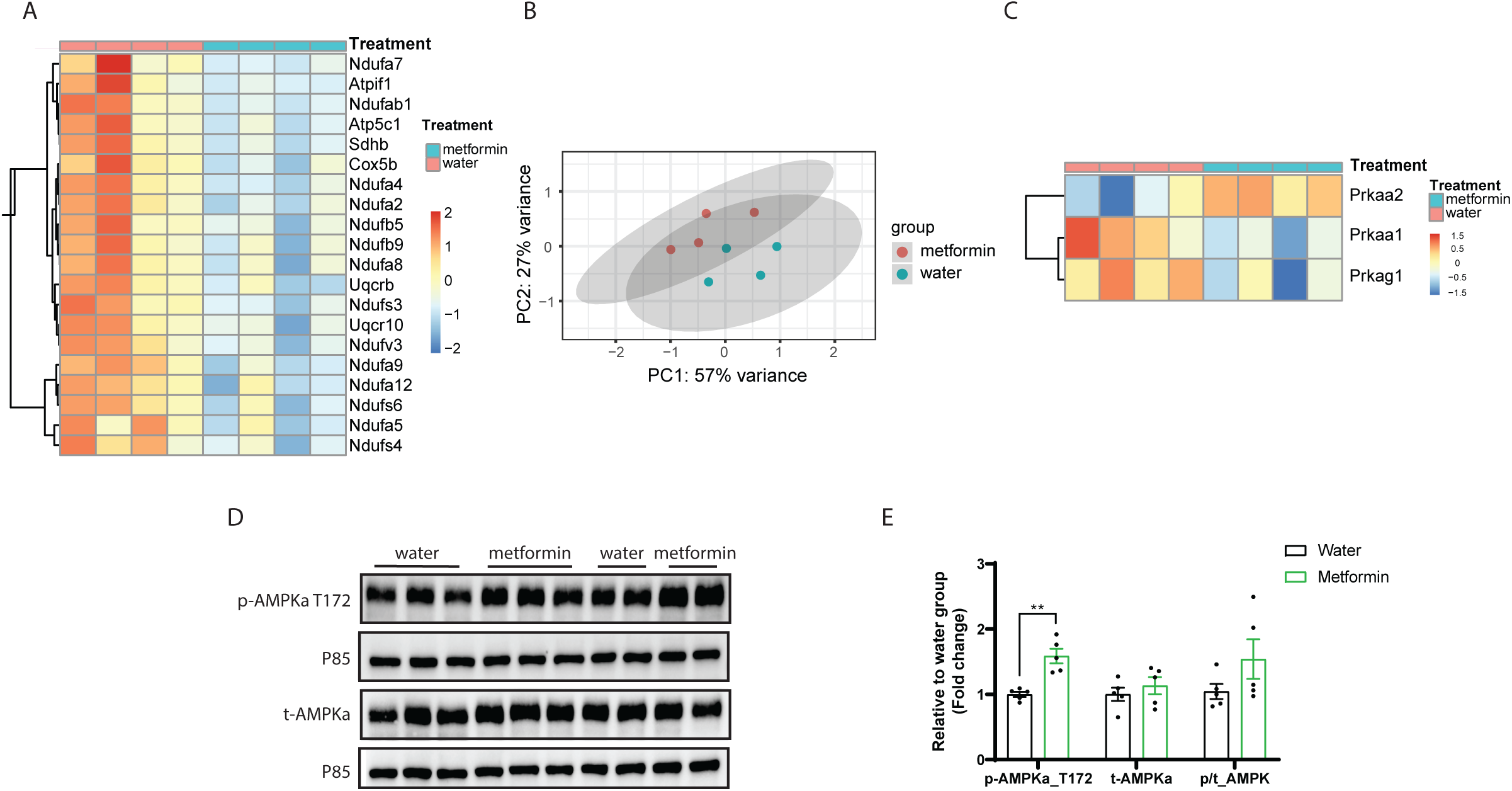
Short-term metformin treatment impacts mitochondrial components and activates AMPK signaling. (A) Metformin reduces protein levels of components of the electron transport chain in intestinal tissue. (B) Principal component analysis using proteins annotated in the AMPK signaling pathway (KEGG: mmu04152). (C-D) Intestinal protein abundance in AMPK signaling pathway in water versus metformin group. (E-F) Western blot and quantification of the expression levels of AMPKα and phospho-AMPKα T172. Analysis within panel A and B was performed with R using data generated from Proteome Discoverer. Data in panel F represent as means ± SEM and paired student-t test was used for statistical analysis. * p<0.05; ** p<0.01; *** p<0.001.

## Discussion

Metformin is a widely prescribed anti-diabetic drug with uncertain target tissues and mechanism of action. Identification of metformin’s target tissue will facilitate the understanding of metformin’s glucose lowering effects and may promote the development of additional therapies for metabolic diseases. Previous studies suggested the potential of the intestine as an important metformin target since metformin concentrates to high levels in intestinal tissue [11; 12]. Intravenous administration of metformin or reducing intestinal metformin absorption impairs metformin’s effects to lower glucose production [42; 43]. Few studies have investigated the role of metformin on intestinal function and its association with systemic glucose homeostasis. Among the studies focusing on the intestine, the beneficial effects of metformin were suggested to closely associated with its role on stimulating intestinal secretion of GLP-1 and GDF15 which are known to regulate systemic glucose homeostasis [27; 37; 44]. To our knowledge, no study has directly assessed metformin’s effects on intestinal metabolic function using stable isotopic tracers. In this study, we assessed intestinal fructose metabolism after an oral fructose bolus (1:1 ratio of unlabeled:^13^C-labeled fructose) to probe intestinal carbohydrate metabolism after short-term metformin treatment. This methodology is based on recent findings from our lab and others that fructose is rapidly metabolized in the small intestine and can be readily used by the intestine to produce glucose [17; 18].

In this study, we observed that 30 minutes after fructose gavage, the majority of glucose within intestinal tissue contained fructose-derived carbons and half contained three fructose-derived carbons (Figure 3A, E, and F) in both metformin treated mice and controls. In systemic circulation, enrichment of glucose with fructose-derived carbon was ~ 50% lower than in intestinal tissue (Figure 3B vs 3D) indicative of dilution of intestinally derived glucose in systemic circulation. This supports prior work indicating that fructose is readily used as metabolic substrate for glucose production in the intestine and this contributes significantly to whole-body glucose production [17; 18]. Our data demonstrates that short-term metformin treatment reduced intestinal glucose production from fructose and fructose delivery to the liver (Figure 2 and 3).

Metformin reduced intestinal glucose production, and this is associated with enhanced glucose tolerance (Figure 1C and D). Importantly, these beneficial effects occurred prior to significant weight changes (Figure 1A). Whether the effect of metformin to reduce intestinal glucose production is a major mechanism by which metformin enhances systemic glucose tolerance remains to be defined.

With the prevalence of high fructose corn syrup in the typical Western diet, the potential for fructose consumption as a contributor to metabolic disease has attracted increasing attention [45]. Studies conducted in both human and rodents support the hypothesis that excessive fructose consumption can contribute to metabolic disorders including obesity, insulin resistance, and NAFLD [46–48]. Intestinal tissue is a major site of fructose metabolism and shields the liver from fructose induced metabolic disease [17], as hepatic fructose metabolism is essential for the development of fructose related pathologies like increased de novo lipogenesis, steatosis, and insulin resistance [30]. A prior study suggested that fructose-induced metabolic disease may be attenuated by metformin treatment in mice although this was attributed to an effect of metformin to alter the microbiome and protect against deterioration of gut barrier function [49]. Our results are consistent with a protective effect of metformin against fructose-induced metabolic disease as we demonstrated that metformin decreased fructose delivery to the liver (Figure 2E-G). These results might point towards a particularly beneficial effect of metformin in people with diabetes and metabolic disease that consume excessive amounts of dietary sugar.

In addition to decreasing fructose delivery to the liver, we also demonstrated that metformin decreased intestinal fructose metabolism. Specifically, intestinal F1P levels were nearly halved in the intestinal tissue of metformin treated animals when compared to controls (Figure 2B). Additionally, significantly lower enrichment of isotope labeling was detected in the downstream three-carbon phosphates and TCA cycle metabolites (Figure 2C and D, Figure 4 A-C). It is interesting that metformin treatment not only decreased intestinal fructose metabolism, but also reduced the amount of fructose delivered to the liver, suggesting the possibility that metformin treatment reduced intestinal fructose absorption. Indeed, unbiased proteomics indicated that the abundance of luminal fructose transporter GLUT5 and the basolateral glucose/fructose transporter GLUT2 were both reduced by metformin treatment (Figure 5F). This finding seems contradictory to a recent study where metformin increased GLUT2 expression in the colon and to a lesser extent in the ileum [13]. This discrepancy may be due in part to differences in intestinal segments analyzed between the two studies. Further investigation will be required to determine the impact of metformin on distinct intestinal segments.

Gastrointestinal symptoms including bloating and diarrhea are the most prevalent side effects of metformin treatment and these symptoms are often experienced in the first few weeks following treatment initiation [50]. It is also well-established that defects in intestinal fructose metabolism can cause significant GI disturbances [51–54]. It is interesting to speculate that the ability of metformin to impair fructose absorption may contribute to its unwanted gastrointestinal effects. Future studies may include investigation into how metformin alters intestinal epithelial absorptive function and substrate transport.

Most studies investigating metformin’s mechanism of action focus on specific putative targets include Complex 1, AMPK, mitochondrial glycerophosphate dehydrogenase, PEN2, and others [14; 40; 49; 55; 56]. Unexpectedly, our proteomics data conducted on metformin treated intestinal tissue suggests that metformin produces broad-based reductions in the levels of many metabolic enzymes including key enzymes in glycolytic, gluconeogenic, and fructolytic pathways. These results are consistent with our conclusions regarding metformin’s multiple metabolic effects derived from isotope tracing data that fructose metabolism, glucose production, and other aspects of intestinal carbohydrate metabolism are all significantly impacted by metformin treatment. This novel finding extends our knowledge of metformin’s potential mechanistic effects and points to novel mechanisms of action.

Broad-based effects of metformin on the levels of components of the mitochondrial electron transport chain are also indicated by the proteomics data analysis. Metformin is known to regulate mitochondrial function by inhibiting Complex I of the mitochondrial electron transport chain [40]. However, the physiological relevance of this mechanism is debated as this mechanism requires high concentrations of metformin (~ 150 µM) within the mitochondrial matrix [32]. At the therapeutically relevant doses that we used in this study, we observed that metformin reduced citrate levels and increased malate levels in the portal circulation (Figure 4G and H). This suggests inhibition of malate dehydrogenase and Complex I of the electron transport chain as malate dehydrogenase activity is coupled to Complex I activity. However, as the proteomics analysis indicated that the expression levels of many Complex I subunits including multiple Ndufa, Ndufb and Ndufs subunits were significantly down-regulated by metformin treatment (Figure 6A), we cannot determine whether metformin’s apparent effect to inhibit Complex I activity might be mediated by direct inhibition or through metformin’s more general effect to reduce levels of Complex I components.

AMPK activation is often invoked as a key mechanism mediating metformin action [7; 36]. Although, recent studies suggest metformin relies on both AMPK-dependent and AMPK-independent mechanisms to mediate its beneficial effects [9]. We also observed that the intestinal AMPK signaling was impacted by metformin treatment (Figure 6C and D). Metformin increased phosphorylation of AMPK at Thr172 potentially consistent with increased AMP/ATP ratio (Figure 6E and F). Again, because of the broad-based effects of metformin on metabolic and mitochondrial protein levels, we cannot readily define the mechanism by which metformin treatment leads to AMPK activation. Neither could we determine whether AMPK signaling participates in metformin’s effects to reduce intestinal fructose metabolism and glucose production.

This study supports our recent work indicating that the intestine is a major source of circulating TCA cycle intermediates [31], and that measurement of fructose-derived label in portal and possibly peripheral circulating blood TCA cycle metabolites may provide a useful index of intestinal metabolic function. In the future this approach may be useful for assessing intestinal metabolic function in large-scale human studies and has the potential to open new avenues for investigating intestinal metabolic function in a wide range of scenarios. In this study, we used this approach to demonstrate that the intestine is a target for therapeutic doses of metformin and short-term metformin treatment impacts intestinal glucose and fructose metabolism. Moreover, proteomics analysis suggests that metformin reduces the intestinal levels of a broad array of metabolic enzymes and transporters involved in sugar metabolism. Additional work will be required to define mechanisms by which metformin reduces the levels metabolic enzymes and impacts key metabolic pathways within the intestine to enhance glucose homeostasis.

## Author Contributions

**MAH, WT, SAH, AS and IA** designed, performed, and interpreted the mouse experiments. **GZ, WT, MAH, and SAH** designed, performed, and interpreted metabolomics analysis. **PAG, MAH, IA, and WT** designed, performed, and interpreted proteomics analysis. **MAH** conceived of, designed, and supervised the experimental plan, and interpreted experiments. **WT** and **MAH** wrote the manuscript. All authors edited the manuscript.

## Acknowledgements

This work was supported by NIH grant 5R01 DK121710 (MAH). Metabolomics support was provided by the North Carolina Diabetes Research Center (NCDRC) (P30 DK124723).

## Notes

### Competing Interest Statement

Mark A. Herman has received research support from Eli Lilly and Co.

### Summary of Updates

Updated figures and corresponding text.

